## Supplementary material for "Fine tuning of calcium constitutive entry by optogenetically-controlled membrane polarization: impact on cell migration": Figure 3 supplement

### Supplementary data

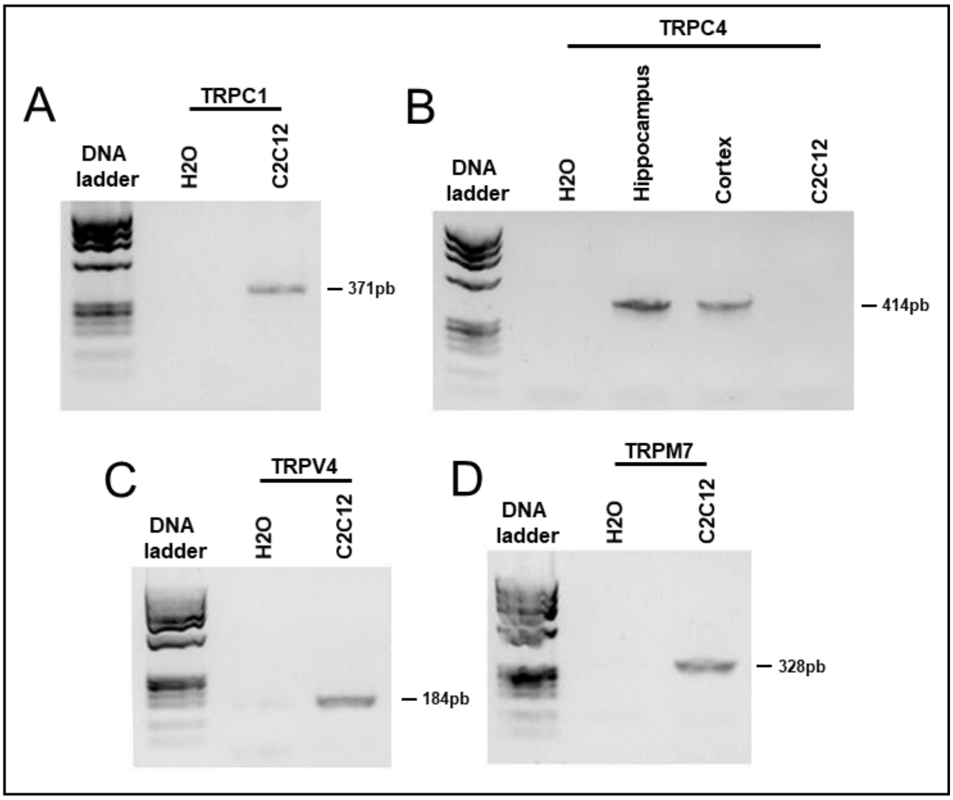


**Figure 3-figure supplement 1**. Expression of different TRP channel mRNAs in C2C12 myoblasts using standard RT-PCR. Reverse-transcribed RNA (150 ng) from C2C12 myoblasts was added to the PCR mixture and PCR products amplified in 35 cycles. PCR products were separated on agarose gels and stained with ethidium bromide. Water served as negative control. (**A**) TRPC1. (**B**), TRPC4. Hippocampus and cortex served as positive control. (**C**) TRPV4. (**D**) TRPM7.
